# Housekeeping Gene Expression Normalization in Transcriptomics Mitigates Data Leakage in Machine Learning Models

**DOI:** 10.64898/2026.04.24.720637

**Authors:** Guilherme Taborda Ribas, Cristian Vidal Riella, Dieval Guizelini, Maurício Menegatti Rigo, Leonardo Vidal Riella, Thiago De Jesus Borges

**Affiliations:** Center for Transplantation Sciences, Massachusetts General Hospital/Harvard Medical School, 149 13th st, Boston, 02129, MA, USA; Professional and Technological Education Sector, Federal University of Paraná, R. Dr. Alcides Vieira Arcoverde, 1225, Curitiba, 81520-260, PR, Brazil; Nephrology Division, Department of Medicine, Beth Israel Deaconess Medical Center, 330 Brookline Ave, Boston, 02215, MA, USA; Center for Discovery and Innovation, Hackensack Meridian Health, 111 Ideation Way, Nutley, 07110, NJ, USA

**Keywords:** Data leakage, machine learning, normalization, housekeeping genes, kidney transplantation, transcriptomics

## Abstract

**Background:** Inappropriate normalization can lead to data leakage and overfitting in machine learning models. Accurately identifying housekeeping genes (HKGs) can overcome this problem and is crucial for normalizing gene expression data, particularly in RNA-Seq experiments.

**Results:** First, we demonstrate that the gene expression of commonly used HKGs significantly changes over time due to immunosuppressive treatments in transplant recipients. Using large public transcriptomic datasets of kidney transplantation, we developed a pipeline based on the gene’s coefficient of variation, stability, and Gini coefficient, and identified nine stable and better-suitable HKG candidates. Our results demonstrate that these HKGs improve the robustness and generalizability of machine learning models by minimizing data leakage, as evidenced by superior performance compared to benchmark methods like median ratio normalization and trimmed mean of M values.

**Conclusions:** This approach enables more accurate comparison of gene expression datasets across different clinical scenarios, improving the reliability of biomarker identification and enhancing personalized treatment strategies.

## 1 Background

To calculate the differential expression of genes, discover biomarkers, and model machine learning predictors, it is fundamental to normalize the data expression using baseline levels of control genes. The current benchmark methods to normalize transcript expression data in RNA-Seq experiments are median ratio normalization (MRN) and trimmed mean of M values (TMM), included in packages DESeq2 [1] and EdgeR [2], respectively. Those methods are based on cross-sample normalizations [3][4][5], which can be a source of data leakage and following overfitted machine learning models [6][7], compromising the reproducibility and generalizations of the models on new datasets [8]. An alternative to cross-sample normalization is defining housekeeping genes (HKGs) for a specific niche, such as kidney transplanted recipients, and normalizing each sample by them independently of other samples.

Although the biological definition of housekeeping genes is still not fully determined, one can define them as genes with constant expression in all conditions, with an essential role in cellular maintenance and a conserved sequence in evolutionary history [9][10][11]. Nevertheless, since the expression levels of a gene can vary with drug treatment [12] and clinical conditions [13][14][15][16], different diseases and treatments can lead to a different set of HKGs. Thus, they should be defined for cohorts subject to specific treatments and conditions and be carefully analyzed before their application in experiments and diagnosis.

Transplantation is the treatment of choice for patients with end-stage kidney disease. In this context, it is essential to define housekeeping genes to support biomarkers discovery and to build tools based on RNA-Seq to indicate potential diagnosis of rejection. In general, kidney transplant recipients are treated with immunosuppressive medications to avoid rejection by the immune system. Some of them can affect some crucial cell maintenance processes, such as DNA repair, leading to alterations in cell physiology and the expression of some genes, including traditional housekeeping genes [17][18]. The commonly used HKGs *GAPDH, ACTB*, and *B2M* have their expression dysregulated in the biopsy of kidney transplanted recipients [19], underlying the need for a better definition of housekeeping genes in this population in different tissues, including blood.

To address the lack of information about appropriate HKGs to serve as controls in experiments using peripheral blood of kidney transplanted recipients, we initially develop a workflow based on large public datasets of RNA-Seq and RNA microarrays with different time points and clinical outcomes (Table 1). We explore well-established methods to define non-differential expression genes. We then apply statistical concepts like coefficient of variance, Gini coefficient, and pairwise stability in an extensive dataset and leverage machine learning concepts to find the best HKG candidates. Moreover, we investigate their conservation across vertebrate species and explore their biological functions and processes. Importantly, all the findings in RNA-Seq were validated in RNA microarrays, demonstrating our method’s consistency and ability to generalize the findings across multiple kidney transplant datasets.

**Table 1.**
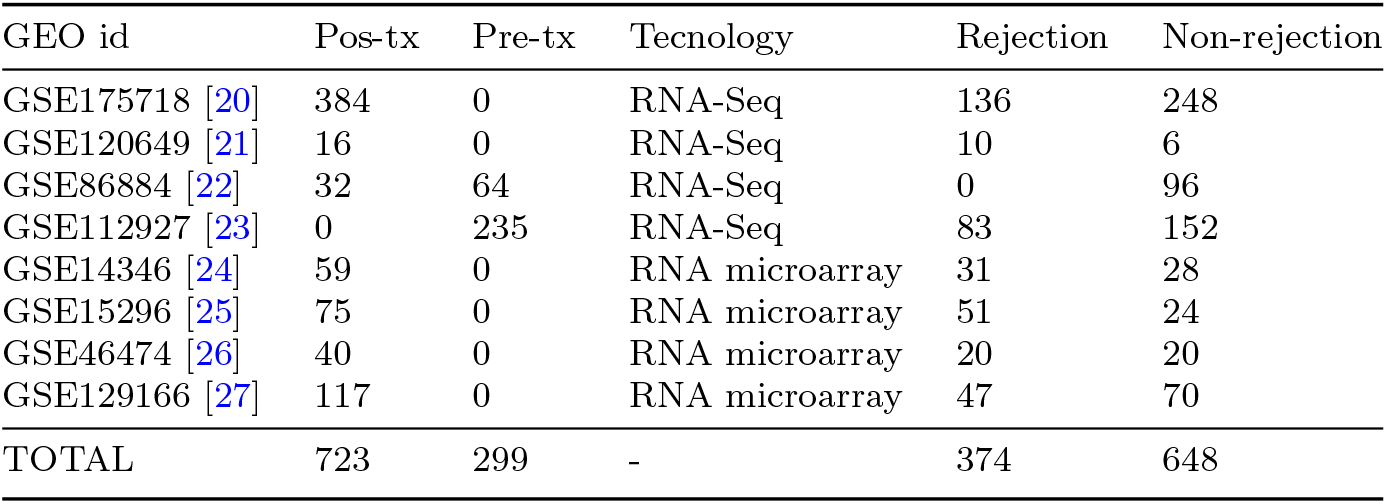
Public peripheral blood transcriptomics data deposited in NCBI.

| GEO id | Pos-tx | Pre-tx | Tecnology | Rejection | Non-rejection |
| --- | --- | --- | --- | --- | --- |
| GSE175718 [20] | 384 | 0 | RNA-Seq | 136 | 248 |
| GSE120649 [21] | 16 | 0 | RNA-Seq | 10 | 6 |
| GSE86884 [22] | 32 | 64 | RNA-Seq | 0 | 96 |
| GSE112927 [23] | 0 | 235 | RNA-Seq | 83 | 152 |
| GSE14346 [24] | 59 | 0 | RNA microarray | 31 | 28 |
| GSE15296 [25] | 75 | 0 | RNA microarray | 51 | 24 |
| GSE46474 [26] | 40 | 0 | RNA microarray | 20 | 20 |
| GSE129166 [27] | 117 | 0 | RNA microarray | 47 | 70 |
| TOTAL | 723 | 299 | - | 374 | 648 |

Finally, we demonstrate that the normalization by housekeeping genes found using our method minimizes data leakage during cross-validation in machine learning modeling due to its independence of cross-sample information. It can also improve the area under the ROC curve compared with cross-sample normalizations, MRN, and TMM, even in independent sample normalization (TPM) comparison. The housekeeping gene mining process is organized and documented in a Python package and can be easily used in other datasets.

## 2 Methods

### 2.1 Public NGS RNA-Seq processing

We reprocess 731 samples from three NGS RNA-seq bulk studies from the peripheral blood of kidney transplanted patients. We guarantee that all samples were submitted to the same pipeline and the same reference genome, preserving the reproducibility of the in silico experiment and controlling for the preprocessing confounders. The GEO ID and summary of the samples are in Table 1.

All the raw fastq files are downloaded using sra-toolkit 3.0.0 [28]. We use Fastqc 0.11.9 with standard parameters [29] to evaluate the quality of reads. We trimmed 3 the adapters with Fastp 0.22.0 [30] and quantified the abundance of transcripts with Salmon 1.9.0 [31]. We used the human transcriptome and the annotation named GRCh38.p14 v.44:2023-03-01 from Gencode as transcriptome reference [32].

We exclude pseudogenes from this analysis to keep only transcripts that could be found in microarray. To guarantee the experiment’s reproducibility, we developed a Snakemake [33] workflow to run this entire preprocessing step.

The dataset GSE86884 [22] is a longitudinal study with 96 samples and four time points of collected data: pretransplant, one week, 3, and 6 months after transplantation. None of the patients rejected the transplant at the time of collection. The dataset GSE120649 [21] contains 16 samples where six patients had stable graft function (non-rejection), 6 with antibody-mediated rejection (ABMR), and 4 with T-cell mediated rejection (TCMR). The dataset GSE175718 [20] contains 384 samples where 248 patients didn’t present rejection, 86 presented ABMR, 68 presented TCMR, and 18 had concomitant ABMR and TCMR. The dataset GSE112927 [23] contains 235 pretransplant samples with 152 non-rejection and 83 patients with rejection.

### 2.2 Public microarray processing

We download the CEL files of the following microarray studies based on the peripheral blood of transplanted recipients GSE14346, GSE15296, GSE46474, and GSE129166 (Table 1). The study GSE14346 contains samples from peripheral blood leukocytes, which were excluded, and samples from peripheral blood, where 28 didn’t reject and 31 presented acute rejection. The study GSE15296 contains 51 patients with rejection and 24 non-rejections. The study GSE46474 classifies patients in acute rejection (n = 20) and non-rejection (n = 20). The study GSE129166 contains 70 non-rejection samples and 47 acute rejections, where 21 were diagnosed with ABMR, 17 with TCMR, and 9 with both types of rejection. Only the last study provides ABMR and TCMR classification. All studies were run on Affymetrix Human Genome U133 Plus 2.0 Array platform. We extract the expression levels with the R 4.4.0 package affy 1.82.0 [34].

The metadata was extracted from an *in-home* Python 3.12.1 algorithm. We transformed the names of the probes to symbol genes based on the same reference genome used in NGS RNA-seq processing. To update the probes to that genome, we used MyGene.py 3.2.2 [35] and grouped the duplicated gene symbols by the median of each sample. MyGene package didn’t translate the probes id to genes *MT-CO1* and *UBNX6*; we manually searched for their probes codes on the GEO platform database (https://www.ncbi.nlm.nih.gov/geo/query/acc.cgi?acc=GPL570).

### 2.3 Statistical Analysis

#### Discovering

We use Tximport 1.32.0 [36] to import the abundance files to analyze it with DESeq2 1.44.0 [1]. We test the alternative hypothesis as “greater than” and log2FoldChange equal 0 for Fig 1A, C. In Fig 2A-C, we use an alternative hypothesis, “less than,” and log2FoldChange equals 0.5. The genes of interest have differential expression between pre-transplant vs. one week, 3 months, and 6 months in maximum +0.5 and minimum −0.5 of log2FoldChange. We consider an adjusted p-value to ≤ 0.05 as a significant level. For each study, we use the normalized expression counts from DESeq2 and calculate the coefficient of variation (CV) (equation 1), coefficient of variation of pairwise stability (cvSTB) (equation 2), and Gini coefficient (equation 3) for all *n* genes of *m* samples with *in-home* Python 3.12.1 script.

**Fig. 1.**
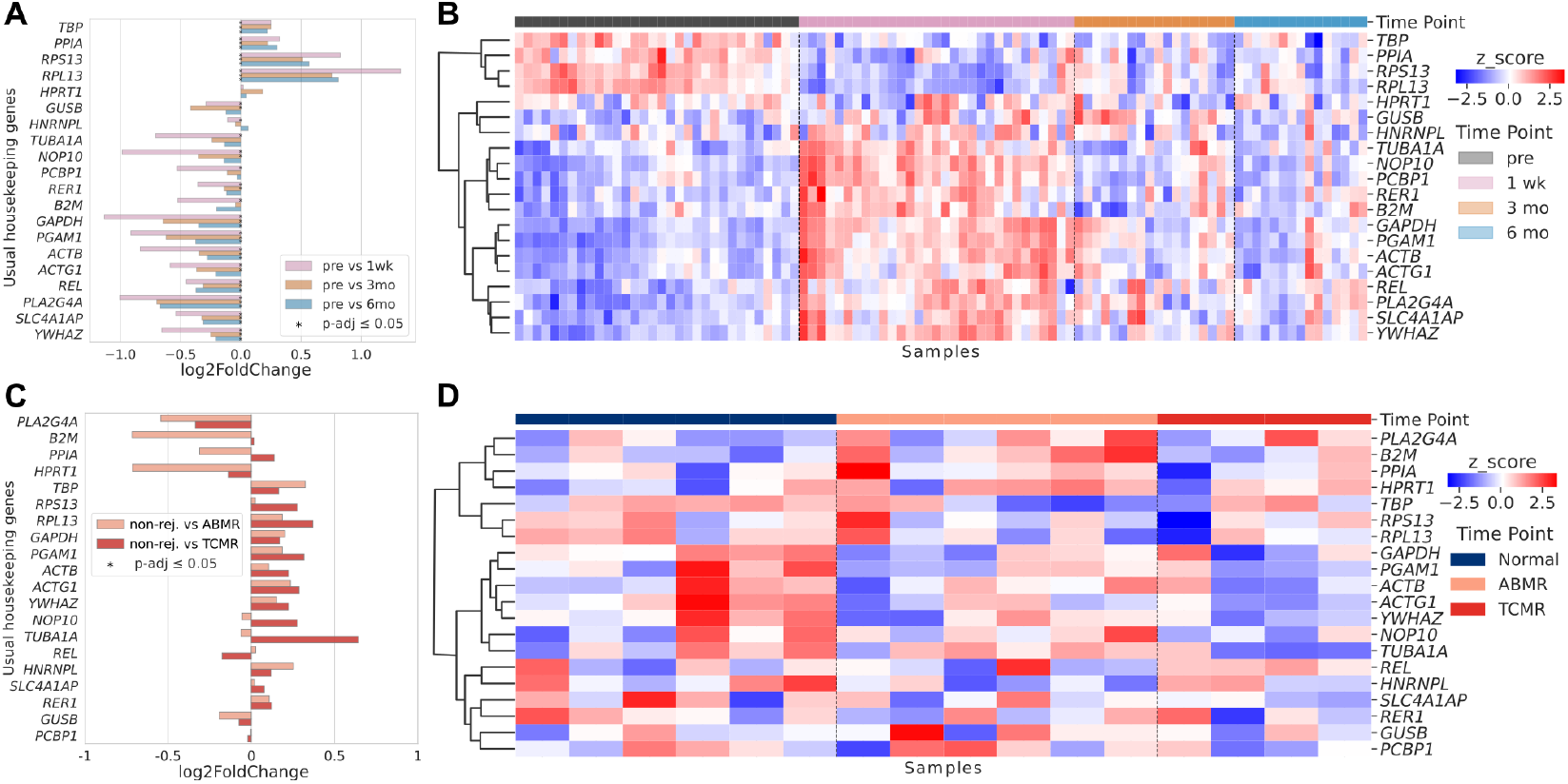
Commonly used housekeeping genes vary in expression according to the patient’s clinical outcome and overtime point in transplantation. (A) Differential expression log2FoldChange and (B) Heatmap of normalized z-score expression of usual HKGs in non-rejection transplanted recipients in different time points. All listed genes have a significant difference (adjusted p-value ≤ 0.05) at least at one time point. (C) Differential expression log2FoldChange and (D) Heatmap of normalized z-score expression of usual HKG in non-rejection, ABMR, and TCMR of transplanted patients. We used DESeq2 to perform differential expression analysis and the statistical significance. To cluster the HKGs on a heatmap, we use average linkage based on Euclidean distance. ABMR: antibody-mediated rejection; TCMR: T-cell-mediated rejection

**Fig. 2.**
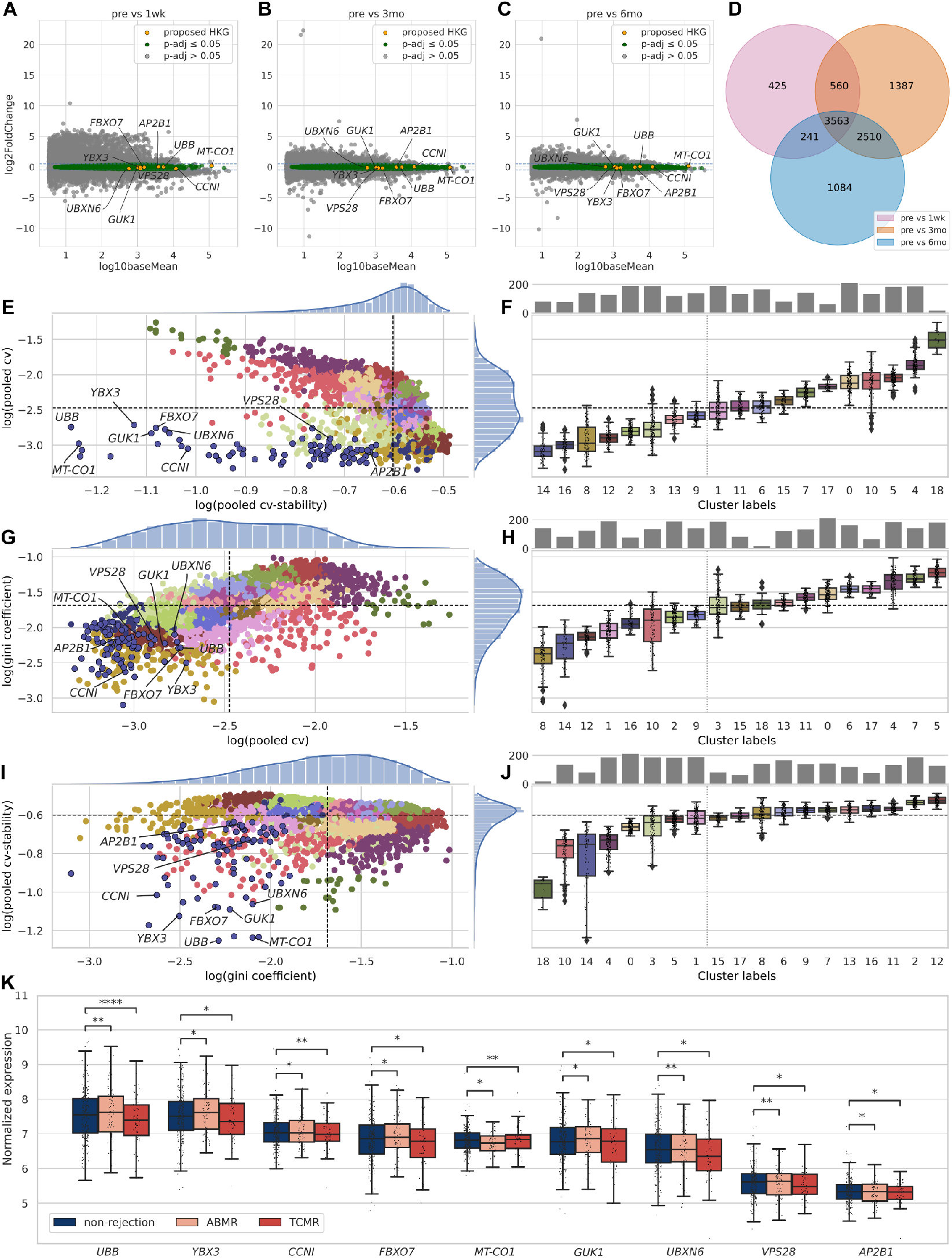
Proposed housekeeping genes present stable expression over all transplanted patients. Gene expressions are in between −0.5 and +0.5 of log2FoldChange in transplanted patients when compared to pre-transplant with different time points, including (A) one week, (B) 3 months, and (C) 6 months after transplantation. (D) Venn diagram showing the intersection of 3,563 genes with log2FoldChange between −0.5 and +0.5 in all time points with adjusted p-value ≤ 0.05. For the 3,563 intersection genes, we display (E) a Scatter plot of the coefficient of variation and cv-stability; (F) a Boxplot of the coefficient of variation of genes clusters; (G) a Scatter plot of the Gini coefficient and coefficient of variation; (H) Boxplot of cv-stability of genes clusters; (I) Scatter plot of cv-stability and Gini coefficient; (J) Boxplot of cv-stability of genes clusters. The dotted lines are the medians of each axis; (K) Two-One-Sided-Test comparing equivalence of expression distributions between non-rejection, ABMR, and TCMR with Brunner-Munzel test considering a Cohen’s d effect size at maximum 0.3. The asterisks indicate statistical significance: * 0.01 *<* p 0.05, ** ≤ 0.001 *<* p 0.01, *** ≤ 0.0001 *<* p 0.001 and ****: p ≤ 0.00001. ABMR: antibody-mediated rejection; TCMR: T-cell-mediated rejection

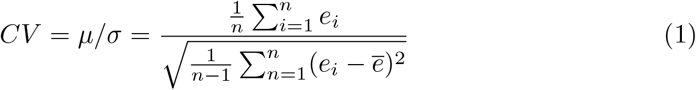

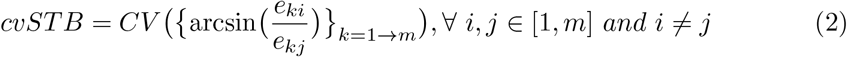

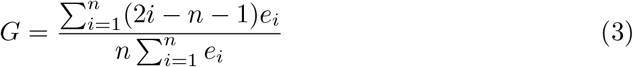

Where, *e* is the gene expression value, *µ* is the average of gene expression and *σ* is the standard deviation,

Since most of the public dataset does not inform batch sample process, demography, and health conditions of the patients, we cluster the samples of each study with a Louvain algorithm from scikit-network 0.31.0 python package [37], we calculate CV and cvSTB for each group and pool the values giving the same weight for each cluster. After calculating the metrics for all genes, we clustered the genes to group those with similar patterns using the Louvain algorithm. We tested if the best candidates have different distributions with the Kruskal-Wallis H-test and performed an equivalent Brunner-Munzel One-Two-Sided-Test (TOST) with a Cohen’s d equal 0.3. Kruskal-Wallis H-test is calculated with SciPy 1.11.4 [38], while TOST is calculated with an *in-home* algorithm.

#### Microarray validation

We calculate the coefficient of variation, coefficient of variance of stability, and Gini coefficient of all genes in each microarray study. We used UMAP from umap-learn 0.5.5 python package [39] to visualize the samples of the nine housekeeping genes in two dimensions. We calculate the K-means clustering for k=2 with scikit-learning 1.4.0. We calculate the Shannon entropy for each cluster with SciPy 1.11.4 [38].

#### Pathway Analysis

We used GSEApy 1.1.1 [40] to filter pathways related to the housekeeping genes. We performed the enrichment analysis against GO Molecular Function and GO Biological Process 2023.

#### Orthology Analisys

We search for orthologues in the HomoloGene database from NCBI. We identify unique Phylum, Classe, Order, Family, Genus, and Species with an in-home script. We download all the vertebrate amino acid sequences with NCBI Datasets tools 16.22.1 [41] and align them using the Clustal Omega 1.2.4 algorithm [42]. For each site of each aligned gene, we calculate the normalized Shannon entropy with SciPy 1.11.4 [38] to quantify the conservation of the amino acids given the homo sapiens sequence as a reference.

#### Machine learning to evaluate normalizations

We perform the analysis on samples of GSE175718 [20]. We detect outliers with HDBSCAN 0.8.36 python package [43] on non-cross-sample normalization TPM. We calculate the pairwise correlation distance between genes and keep only one gene from groups of highly correlated genes. We calculate the pairwise R Pearson correlation between the 9 housekeeping genes, we cluster them based on distance correlation and average hierarchical linkage method. We chose a cophenetic distance for 4 cluster based on silhoutte, Calinski Harabasz and Davies Bouldin scores (FIG S3). We used SciPy 1.11.4 and scikit-learning 1.4.0 to perform all the previous analysis.

We calculate the original MRN and TMM normalizations based on the package conorm (https://pypi.org/project/conorm/), as well as create two python classes to integrate an adapted version of MRN and TMM transformations to a scikit-learning pipeline [45]. To calculate the normalization factors based on different means, we used follow equations for each sample. Where *e* is the expression values of *n* genes used as housekeeping genes.

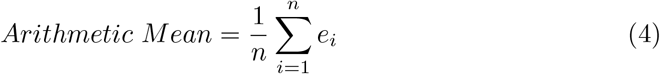

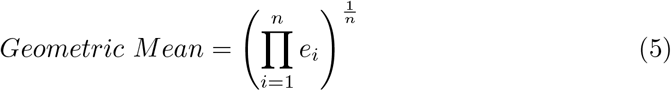

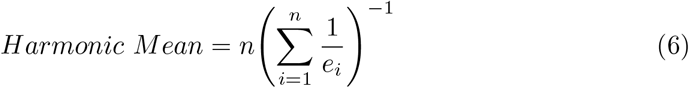

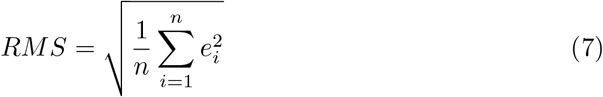

To normalize the expression dataset, we divided each gene expression value by the normalization factors calculated by sample (equation 8). Where *e*_*k*_ is a gene expression value in each *m* sample.

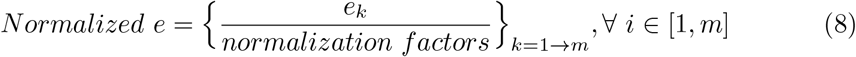

To avoid imbalanced classes, we used the method RandomUnderSampler from imbalanced-learn 0.12.0 Python package [46]. We construct a pipeline in scikit-learning 1.4.0 [45] to undersampling, select features, and train/test a Random Forest for different normalization methods for the same set of subsamples. We apply crossvalidation and calculate the AUC for each set of training and test. The results for all normalization factors are in Fig A4.

#### Data manipulation and plotting

We use Pandas 1.5.3 [47], anndata 0.10.5 [48] and Numpy 1.26.4 [49] to manipulate the data and Matplotlib 3.8.0 [50], Seaborn 0.11.2 [51] and upsetplot [52] to plot them.

The entire coding process is available online in Harvard Dataverse, including Snake-make workflows, bash scripts, notebooks, and a new Python package we developed during this investigation.

## 3 Results

### 3.1 Gene expression of commonly used housekeeping genes varies in transplanted patients

We assess whether commonly used HKGs [19] keep constant expression levels in peripheral blood from kidney transplant recipients at different time points (one week, three months, and six months after transplantation) using the public dataset GSE86884 [22]. Most genes are differentially expressed at least one time point in post-transplant when compared to a pre-transplant baseline (Fig 1A). The first-week post-transplant shows a bigger absolute change for most housekeeping genes. For example, *GAPDH* expression is downregulated by a log2FoldChange of −1.1 at one-week post-transplant compared to pre-transplant levels. Six months post-transplant, it is downregulated by a log2FoldChange of −0.3, indicating differences in its expression dynamics through time (Fig 1A). We also analyzed the normalized expression profile at each time point and found that all analyzed genes varied expression levels at different time points (Fig 1B). Some HKGs (e.g., *TBP, PPIA, RPS13*, and *RPL13*) have a high expression value in pre-transplant, while other genes have high expression levels after transplantation, mainly in the first week and three months (Fig 1B).

Next, we analyze those common HKGs in the context of non-rejection and different types of kidney rejection events: T-cell-mediated rejection (TCMR) and antibody-mediated rejection (ABMR) from a different data set (GSE120649 [21]). None of those genes present a statistically significant difference. However, *B2M* presented a log2FoldChange of −0.7 with a p-adjusted = 0.08 in ABMR; a similar behavior is seen in *HPRT1* in ABMR and *TUBA1A* for TCMR in terms of log2FoldChange absolute values (FIG 1C-D).

Similarly to our results, all those commonly HKGs presented variability through time and different conditions in kidney biopsy of transplanted recipients [19][53], breast cancer [14], brain stroke [13], and diabetes [15]. Our data demonstrate that commonly used HKGs should be validated before being used as reference genes in transcriptional studies involving peripheral blood of kidney transplant recipients, as their expression can be influenced by immunosuppression or surgical stress.

### 3.2 Identification of housekeeping genes in NGS RNA-Seq

To determine potential alternatives for the traditional housekeeping genes, we reprocess and analyze 731 samples of public RNA-Seq from the peripheral blood of kidney transplant recipients (Table 1). We set a pipeline to keep only genes with low variability in time and for different clinical outcomes: pre-transplant, one week, three or six months after transplantation; non-rejection, T-cell-mediated rejection (TCMR) and antibody-mediated rejection (ABMR) with or without donor-specific antibodies (DSA). First, we exclude differentially expressed genes at three different time points, considering pre-transplant as the baseline and controlling for gender, ethnicity, and age. We calculate the non-differential expression genes between pre-transplant samples versus one week (Fig 2A), three months (Fig 2B), and six months (Fig 2C) after transplantation in 96 non-rejection patients from study GSE86884 [22]. We then select the genes that exhibit a log2FoldChange between −0.5 and +0.5 with adjusted p-value ≤ 0.05 across all comparisons: pre-transplant vs. one week, three months, and six months post-transplant. In total, 3,563 genes are considered equivalent at all time points (Fig 2D).

Secondly, since some public studies cannot provide complete patient metadata information, we cluster all samples using a Louvain unsupervised algorithm to create homogeneous subgroups (Fig A1). For all 3,563 genes, we calculate the coefficient of variation (CV) with equation 1, coefficient of variation of pairwise stability (cvSTB) (equation 2), and Gini coefficient (equation 3). The ideal housekeeping genes must have a low CV through all samples, showing that the expression levels are at the same level independently of the patient’s condition. The same is true for cvSTB; the log ratio of all paired gene combinations is calculated to assess how much the expression of one gene varies with another in different conditions [54][55]. Following the same idea, a lower Gini coefficient indicates more stability of the gene expression in all samples [56][57][58].

To select the genes in an unbiased way, we perform another unsupervised cluster on the CV, cvSTB, and Gini per gene (Fig 2E, G, I) and choose the group of genes with the lowest values of those metrics in 635 samples with rejection and non-rejection classes. Group 14 has the best candidates for HKGs since this group has values for the CV, Gini coefficient, and cvSTB lower than the medians of each of these metrics in all 635 samples (Fig 2F, H, J). This strategy solves the known problem of defining arbitrary cutoff levels of HKG expressions [11]. A total of 85 candidates were selected for the following filtering step.

After selecting the genes consistently expressed in all samples, we ask whether some of those 85 targets are differentially expressed by comparing rejection and non-rejection states in 384 posttransplant samples from the GSE175718 [20] study. We performed the Kruskal-Wallis H-test to identify genes with significant differences in expression across non-rejection, ABMR, and TCMR (Table A1). The genes *AKT2, ANKRD11, BTG1, CYLD, EWSR1, FUS*, and *PRRC2C* have p-values ≤ 0.05, indicating their expression levels were significantly different among the outcomes. We exclude them from the HKG candidates list.

Since the previous test doesn’t guarantee equivalence of the 78 remaining candidates, we perform the Two-One-Sided-Test (TOST) with the non-parametric Brunner-Munzel test to verify whether the 78 genes have Cohen’s d effect size less than 0.3 when compare non-rejection vs ABMR and non-rejection vs TCMR (Table A2). Only the genes *AP2B1, CCNI, FBXO7, GUK1, UBB, UBXN6, VPS28, YBX3*, and *MT-CO1* have a significant adjusted p-value ≤ 0.05 for this equivalence test (Fig 2K). In sum, nine genes are stable in different conditions and time points, presenting low CV, low cvSTB, and low Gini coefficient, and are statistically equivalent between non-rejection and rejection. *CCNI, GUK1, UBB, UBXN6*, and *VPS28* were also defined as housekeeping genes in another cross-non-disease tissue study [59]. Those selected genes are considered for validation in the Microarray platform.

### 3.3 Validation of housekeeping genes in microarray

To validate the selected housekeeping genes in a microarray platform, we calculate the CV, Gini coefficient, and cvSTB for all genes in four RNA microarray datasets (Table 1). As observed in RNA-Seq, the genes AP2B1, *CCNI, FBXO7, GUK1, UBB, UBXN6, VPS28, YBX3*, and *MT-CO1* have low CV (Fig 3A-D), low Gini coefficient (Fig 3E-H), as well as low cvSTB metric (Fig 3I-L). The gene *UBB* has the lowest value for all metrics in all studies, followed by the gene *MT-CO1*. The gene *AP2B1* has the highest metric values in the microarray studies.

**Fig. 3.**
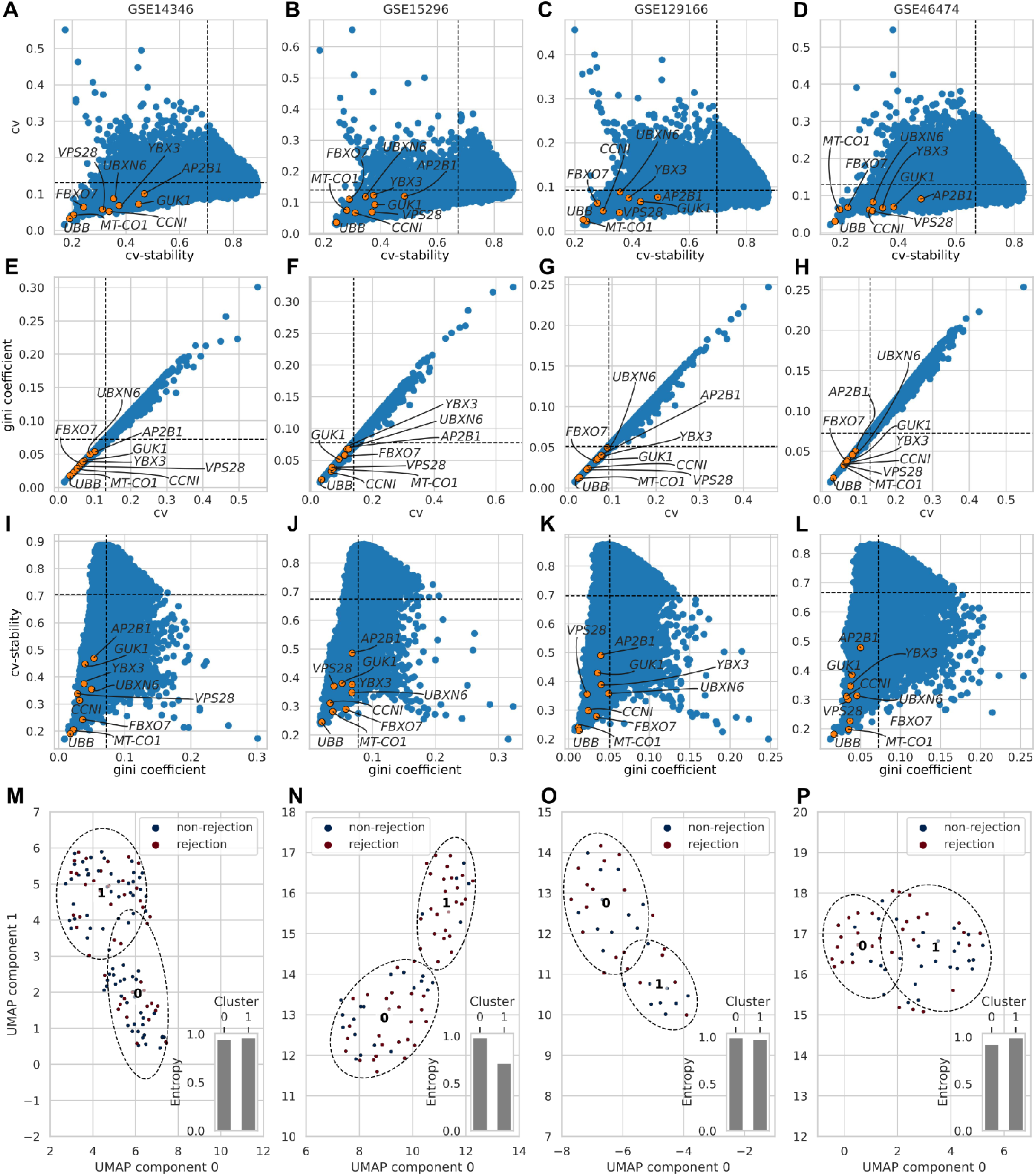
Validation of proposed housekeeping genes in four microarray datasets. (A-D) Coefficient of variation and cv-stability of gene expression in microarray. (E-H) Gini coefficient and coefficient of variation of gene expression. (I-L) cv-stability and Gini coefficient of gene expressions. Yellow points are our proposed HKGs, and all the other genes are in blue. The dotted lines are the axis medians. (M-P) Scatter plot of two UMAP dimensions of proposed housekeeping genes expression clustered in two Kmeans groups and two conditions (rejection and non-rejection). At the right bottom of each scatter is a bar representing the entropy of each Kmeans cluster.

Another way to validate these genes as housekeeping genes is to evaluate how informative they are to distinguish rejection and non-rejection classes in four microarray studies. Genes with low variance cannot distinguish samples between different classes, such as in our example, the classes rejection and non-rejection. To verify, we perform dimensionality reduction and semi-supervised Kmeans clustering (n clusters = 2) using the nine genes and calculate each cluster’s entropy based on each sample’s class. High entropy means high heterogeneity within clusters. Fig 3M-P shows for all microarray datasets that the HKG information does not reduce the entropy of predicted clusters, which is expected of non-informative features [60]. Thus, our results consistently demonstrate that the proposed HKGs have low variability in different clinical outcomes.

### 3.4 Candidate housekeeping genes participate in critical pathways and have highly conserved sites in 630 species of vertebrates

We perform Gene Set Enrichment Analysis [61] for the nine potential housekeeping genes to verify their molecular function and involvement in the biological process. The selected genes are involved in critical biological processes and molecular functions, like maintenance of chromatin structure, DNA repair, stress response, mortality, protein digestion, proliferation, expression and splicing regulation, and cell energy (Fig 4A).

**Fig. 4.**
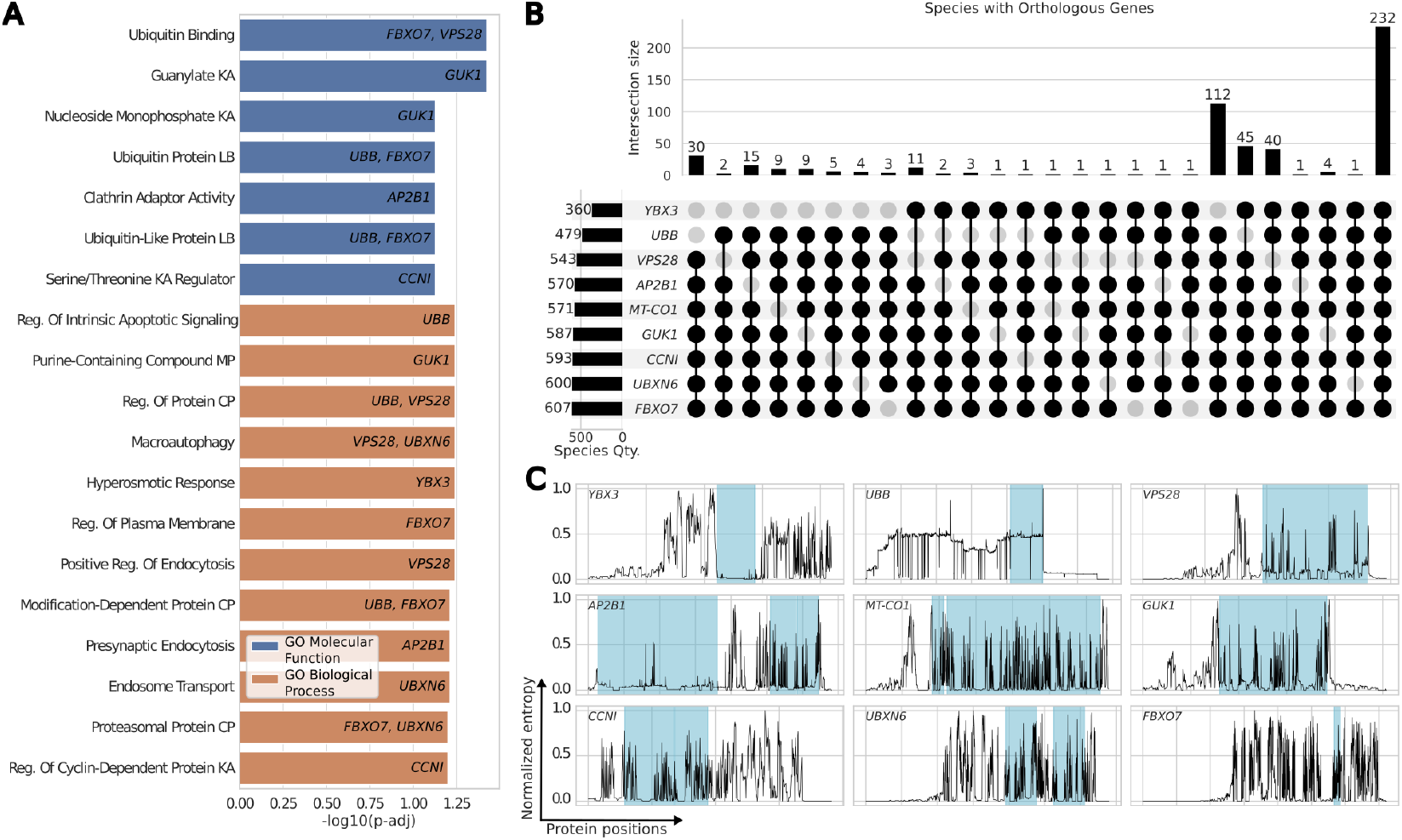
Biological importance and conservation of proposed housekeeping genes. (A) GSEA enrichment analysis for molecular function and biological process of nine housekeeping genes, the vertical axis is the names of the pathways, and the horizontal axis is the statistical significance. (B) Upset plot of species that share at least seven homologous genes. The top barplot shows the number of species sharing the same homologous genes. The left barplot shows the total number of species with the homolog gene. (C) Relative entropy, in the vertical axis, of aligned homologous genes positions (in the horizontal axis) and significant motifs in blue.

An essential characteristic of housekeeping genes is their conservation across species to verify their importance in evolutionary history [9][10]. To evaluate the conservation sites in vertebrates’ homologs, we retrieve the orthologue genes of each proposed housekeeping gene from the HomoloGene Database [62]. In the upset plot in Fig 4B, it is possible to verify the total of species that share orthologues. The gene *FBXO7* is present in 607 species of 630 analyzed, while *YBX3* is present only in 360 species. 536 species share at least seven of the nine housekeeping gene candidates. The upset plots with all combinations for one Phylum, eight Classes, 111 Orders, 288 Families, 500 Genera, and 630 Species are available in Fig A2.

Further, to quantify the conservation of the amino acids given the Homo sapiens sequence as a reference, we perform multiple alignments and calculate the normalized Shannon entropy for each group of homologs. Low normalized entropy values are highly concentrated in essential motifs of each sequence; those regions are highlighted in blue in Fig 4C. For example, in *MT-CO1* between regions 32-69, there is an important Calcium binding region ([63] cd00054), while in *UBB* region 1-228, there is a ubiquitin-like region ([63] cd01803). All regions are described in Table A3.

These data underline their role as housekeeping genes being highly conserved and playing important molecular functions and biological processes, confirming the ability of our method to identify new candidates to be used in non-cross-sample normalization methods.

### 3.5 Normalization by housekeeping genes mitigates data leakage and improves rejection predictor

Finally, to evaluate the impact of the normalization based on housekeeping genes in machine learning modeling, we compare our method with MRN and TMM from the DESeq2 [1] and EdgeR [2] packages, respectively. These methods are the most common and recommended RNA-Seq normalizations in bioinformatics [3][4][5][64]. We also compare HKG normalization with TPM and the inverse hyperbolic sine of TPM. Unlike MRN and TMM, TPM normalizes independently of other samples [3].

After excluding outlier samples flagged by the HDBSCAN algorithm, we define a standard Random Forest model pipeline and perform 25 cross-validations on training and test datasets, ensuring the same conditions for each normalization step. We use the differentially expressed genes identified by Loon V. et al. [20] to select features for the modeling (i.e., *HOMER3, CD14, IFI27, ZEB2, IL18R1, DAAM2, GBP5, NKG7, PATL2, DGKH*, and *SLAMF7*). All training and test dataset classes are balanced with imblearn package [46]. All steps are applied through a scikit-learning [45] pipeline to ensure compliance with best practices in machine learning [65]. We perform the analysis on 378 samples of GSE175718 samples [20].

To normalize the expression data based on our findings, we calculate the pairwise correlation distance [44] between the nine proposed HKGs and cluster them to find groups that could be used as normalization factors (Fig 5A and Fig A3). We use the inverse hyperbolic sine of TPM to reduce the differences in scales between genes and to transform their distribution close to the Gaussian distribution. We calculate the arithmetic mean (equation 4), geometric mean (equation 5), harmonic mean (equation 6), and root means square (equation 7) of the expression levels for all HKG together, of each gene cluster, and pairs within each cluster (Fig 5A). *UBB* and *AP2B1* are used individually as normalization factors since they are not clustered closely to any other gene. After that, we create a new dataset for each normalization factor, subtracting it from all expression values. The normalization calculated based on the arithmetic mean of genes *VPS28* and *MT-CO1* presents the higher AUC metric distribution in the test dataset for the Random Forest model, followed by the harmonic mean of genes *FBXO7* and *CCNI*. The harmonic mean of all proposed housekeeping genes also presents a higher AUC than MRN and TMM.

**Fig. 5.**
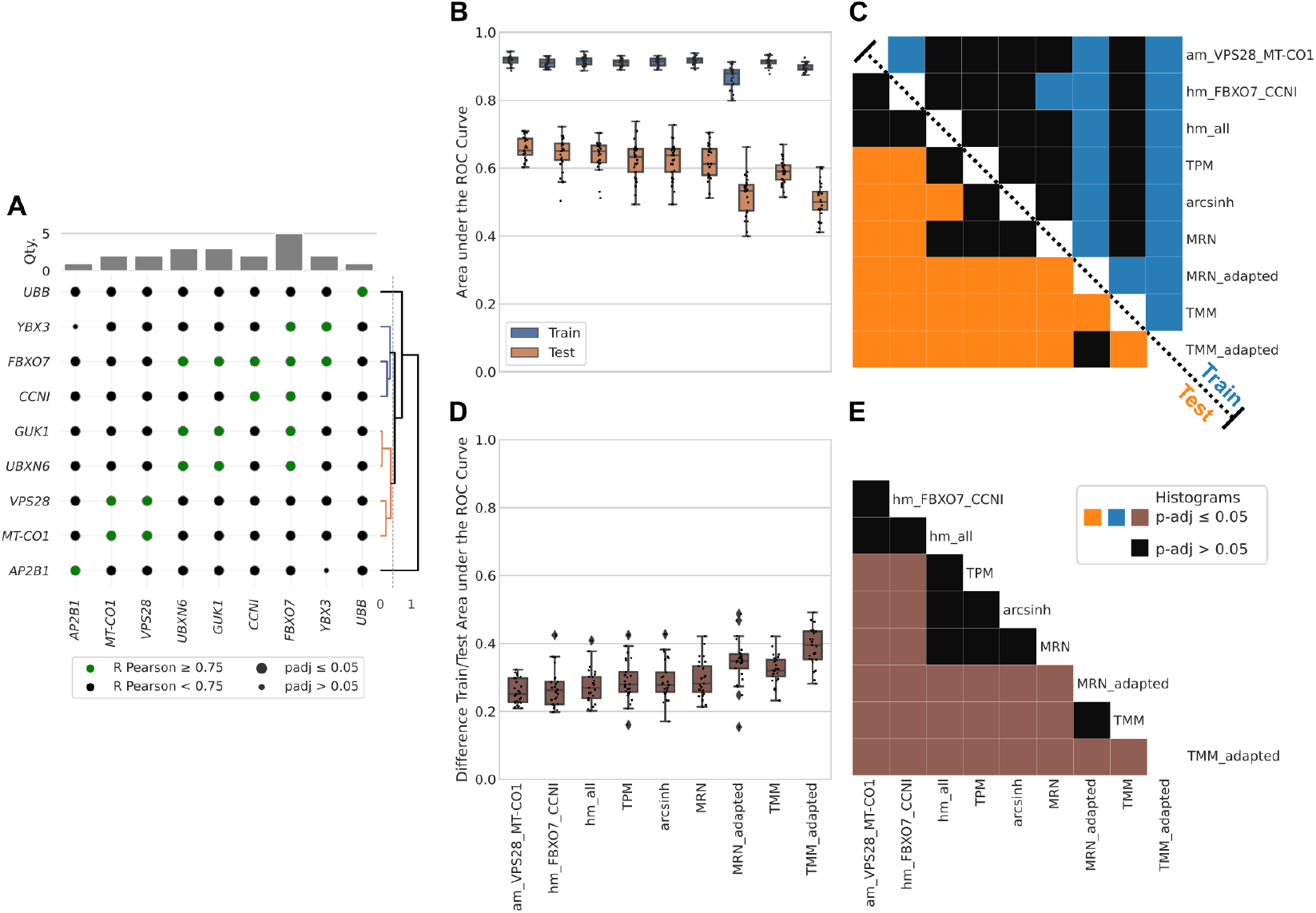
Analysis of data leakage in normalization based on proposed housekeeping genes and benchmark methods. (A) Scatterplot of Pairwise Pearson correlation of proposed housekeeping gene expressions. Color code differentiates the Pearson test’s R ≥ 0.75, and the size differentiates the test significance (adjusted p-value ≤ 0.05). The top barplot is the number of HKGs that are highly pairwise correlated. In the right dendrogram, the hierarchical clustering average for the distance correlation measure correlation forms four groups of normalization factors. (B) Train and test values of AUC metric distributions for different normalization methods for 25 cross-validations. (C) The colored boxes represent statistically significant differences (adjusted p-value ≤ 0.05) in pairwise comparisons of AUC distributions between different normalization methods. The black box represents a non-significant difference. (D) Distributions of differences between train and test AUC values for each normalization method. (E) Adjusted p-values of pairwise comparisons between differences in AUC distributions for different normalization methods. The brown boxes represent statistically significant differences (adjusted p-value ≤ 0.05), while the black boxes represent a non-significant difference.

Since MRN and TMM are cross-sample normalization methods, we test two approaches to assess the impact of data leakage. First, we apply them to the entire dataset before splitting it into training and test sets. We name these approaches MRN and TMM. Secondly, we adapted MRN and TMM to be compatible with the scikit-learning pipeline and minimize data leakage. This way, the training dataset is normalized separately from the test dataset. While these methods calculate scaling factors for each dataset sample, using the scaling factor from training to testing is not possible since they may have different numbers of samples. Therefore, we mitigate the data leakage for MRN and TMM but do not eliminate it. We name these normalizations TMM adapted and MRN adapted.

It is possible to infer that MRN and TMM can deliver an overfitted model because of data leakage, based on Fig 5B. When we mitigate the data leakage using MRN adapted and TMM adapted, we observe the distribution of the area under the ROC curve (AUC) reduce significantly in both the training and test datasets (p-adjusted ≤ 0.05 in Fig 5B). Therefore, they should not be used for modeling predictors but only in investigating differential expressions, in which data leakage is not a problem. Even our versions of MRN adapted and TMM adapted should be avoided. As observed in Fig 5C-D, the differences between AUC distributions from the train to test data set have higher values, showing a reduction of around 0.4 in the AUC median between train and test for the adapted version of TMM and MRN. Conversely, the normalization based on the arithmetic mean of VPS28 and MT-CO1 presented a reduction of around 0.2 (Fig 5C-D). Importantly, our proposed housekeeping genes normalization performed better than both MRN adapted and TMM adapted, delivering a statistically higher AUC in training and testing datasets (Fig 5A-B) and a low AUC difference between train and test results (Fig 5C-D). When we compare the AUC metrics with MRN and TMM, the normalizations based on pairs of genes *VPS28* with *MT-CO1* and *FBXO7* with *CCNI* deliver a better test AUC distribution (Fig 5A-B) and the lowest difference between train and test (Fig 5C-D). Since HKG normalization is a non-cross-sample method, there is no risk of data leakage related to cross-sample normalization. Our method performs better in the test dataset even when we compare housekeeping genes normalization with TPM, which is also a non-cross-sample normalization. This may be related to the technical effect that can be mitigated when housekeeping genes are used in normalization. Since the technical effects affect the entire sample, using the HKGs as normalization factors can reduce the technical differences between samples. We propose that this method can be a simpler solution for batch effect correction for machine learning applications, leading to better predictors.

## 4 Discussion

Previous studies based on the transcriptional profiles of tissue biopsies also showed that extensively used housekeeping genes such as *GAPDH, B2M, RER1, RPL13, TUBA1A*, and *ACTG1* are not reliable reference genes for kidney transplantation patients [19] or kidney conditions [53]. However, until now, no study has defined housekeeping genes specifically for peripheral blood in this population, nor has any study confirmed that traditional housekeeping genes have a high expression variation at different time points or in distinct clinical outcomes (e.g., rejection or non-rejection). Furthermore, differently from other studies [19][53], (i) we investigate the largest dataset of kidney transplanted recipients, (ii) we insert the Gini coefficient along with other metrics to measure stability, (iii) we use unsupervised method to define the stable genes and (iv) we perform the equivalence test (TOST) to make sure that the difference is within a specified interval, while one study considered stable genes the genes that were not differentially expressed [53], which is a mistake, since the absence of differential expression evidence, is not evidence of expression equivalence.

A refined normalization method based on consistently stable genes is essential to create machine learning models to predict rejection responses from peripheral blood transcriptomics with high specificity and sensitivity. We demonstrate that predictors modeled on data normalized by our factors achieve a high area under the ROC curve (AUC) values in the test dataset while effectively avoiding data leakage. Mitigating this source of overfitting is crucial to lead to better models that can contribute to decision-making in health. It can meliorate predictors, leading to minimal invasive diagnostics in liquid biopsy, such as peripheral blood. This approach has the potential to enhance precision and personalized medicine significantly, allowing for earlier diagnosis and continuous monitoring at a low cost to the patients [66][67][68][69].

While our findings are promising, they are currently limited to peripheral blood samples from kidney transplant recipients. Although our methodology can be easily applied to any cohort through the Python package developed in this work, further research is necessary to validate these HKGs across other tissues and in different transplant contexts. Additionally, the lack of demographic data in public RNA-Seq datasets poses a challenge in generalizing these results to broader populations.

Future studies should focus on validating these HKGs in larger, more diverse cohorts, including different demographic groups and conditions. Additionally, exploring the application of these HKGs or the methodology in other organ transplants could provide insights into their broader utility. Investigating the normalization based on these genes in different machine-learning models would also be beneficial, potentially leading to the development of robust, clinically applicable algorithms for early rejection detection.

Developing more reliable normalization methods using niche-specific HKGs marks a significant step in integrating machine learning with clinical genomics. By minimizing data leakage and enhancing model generalizability, our approach could pave the way for more accurate predictors and reproducible biomarker discovery, ultimately contributing to better patient outcomes in transplantation medicine.

## 5 Conclusions

This study presents new normalization factors based on identifying more robust house-keeping genes in transcriptomics datasets to avoid data leakage and mitigate technical differences between samples. Our approach outperforms traditional cross-sample normalization methods such as MRN and TMM, which can deliver overly optimistic results. It also overcomes the TPM, an independent sample method that cannot correct technical artifacts. We screened thousands of genes to construct these normalization factors. We chose the highly expressed ones with low coefficients of variation, high stability, and low Gini coefficient across various conditions and treatments in high-throughput RNA sequencing. Additionally, these genes participate in crucial cellular maintenance processes and are conserved across vertebrate species, adhering to the standard definitions of housekeeping genes.

## Abbreviations

ABMR: Antibody Mediated Rejection
AUC: Area Under the Curve
CV: Coefficient of Variation
DSA: Donor-specific Antibodies
HKG: Housekeeping Genes
MRN: Median Ratio Normalization
ROC: Receiver Operating Characteristic
STB: Pairwise Stability
TCMR: T cell-mediated rejection
TMM: Trimmed Mean of M Values

## Data availability

Supplementary information. All codes and results of *in silico* analysis are deposited in Harvard Dataverse under the DOI https://doi.org/10.7910/DVN/NBR0HG. The Python Package housekeeping-MinerPy version used in this work is deposited in Harvard Dataverse under https://doi.org/10.7910/DVN/PD7KY7. The future updated versions are deposited in GitHub: https://github.com/guilhermetabordaribas/housekeepingMinerPy. The package Documentation can be found at https://housekeepingminerpy.readthedocs.io.

## Declarations

The authors declare no competing interests.

## Appendix A Supplementary Figures and Tabels

**Fig. A1.**
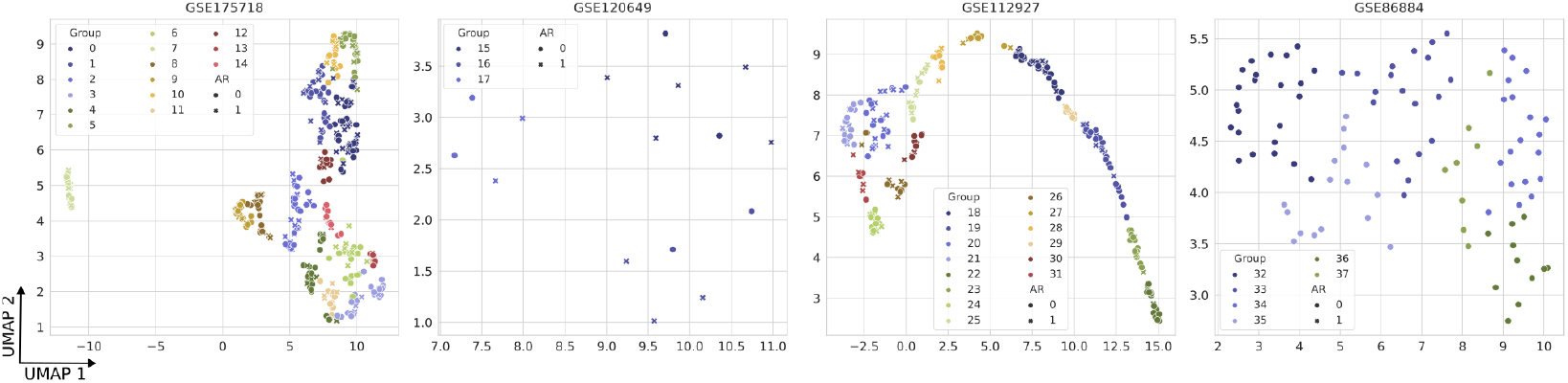
Unsupervised Louvain clustering to group samples due to lack of demographic information. An unsupervised Louvain algorithm creates homogeneous groups for each RNA-seq dataset. The two dimensions of UMAP dimensionality reduction allow the visualizations of groups. The data from each cluster were used to calculate the coefficient of variation (CV), coefficient of variation of pairwise stability (cvSTB), and Gini coefficient.

**Fig. A2.**
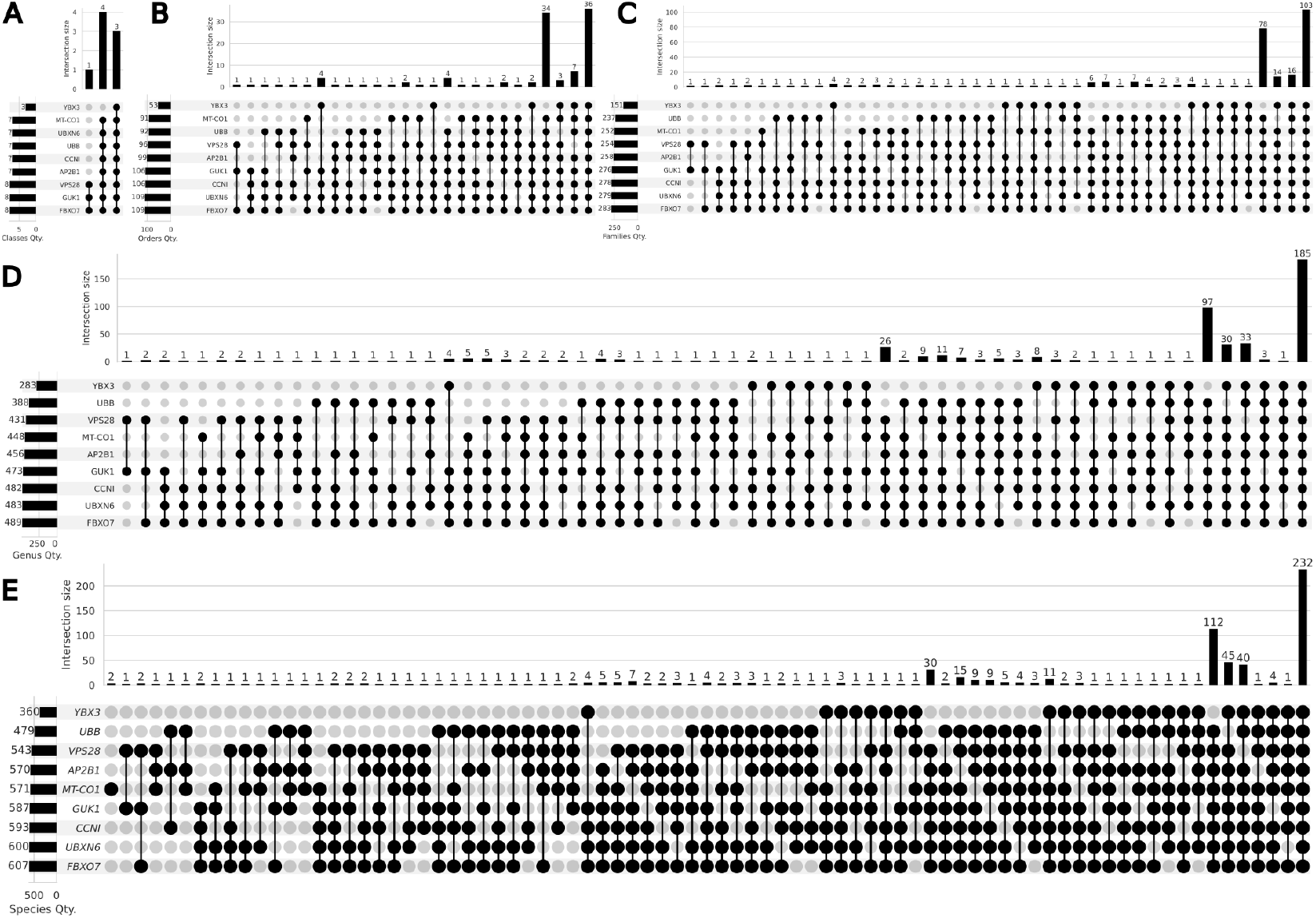
The upset plots with all combinations of homologs for Classes, Orders, Families, Genera, and Species. (A) The upset plot of eight Classes that share homologs. (B) The upset plot of 111 Orders that share homologs. (C) The upset plot of 288 Families that share homologs. (D) The upset plot of 500 Genera that share homologs. (E) The upset plot of 630 Species that share homologs. The top barplot shows the number of species sharing the same homologous genes. The left barplot shows the total number of taxon with the homolog genes.

**Fig. A3.**
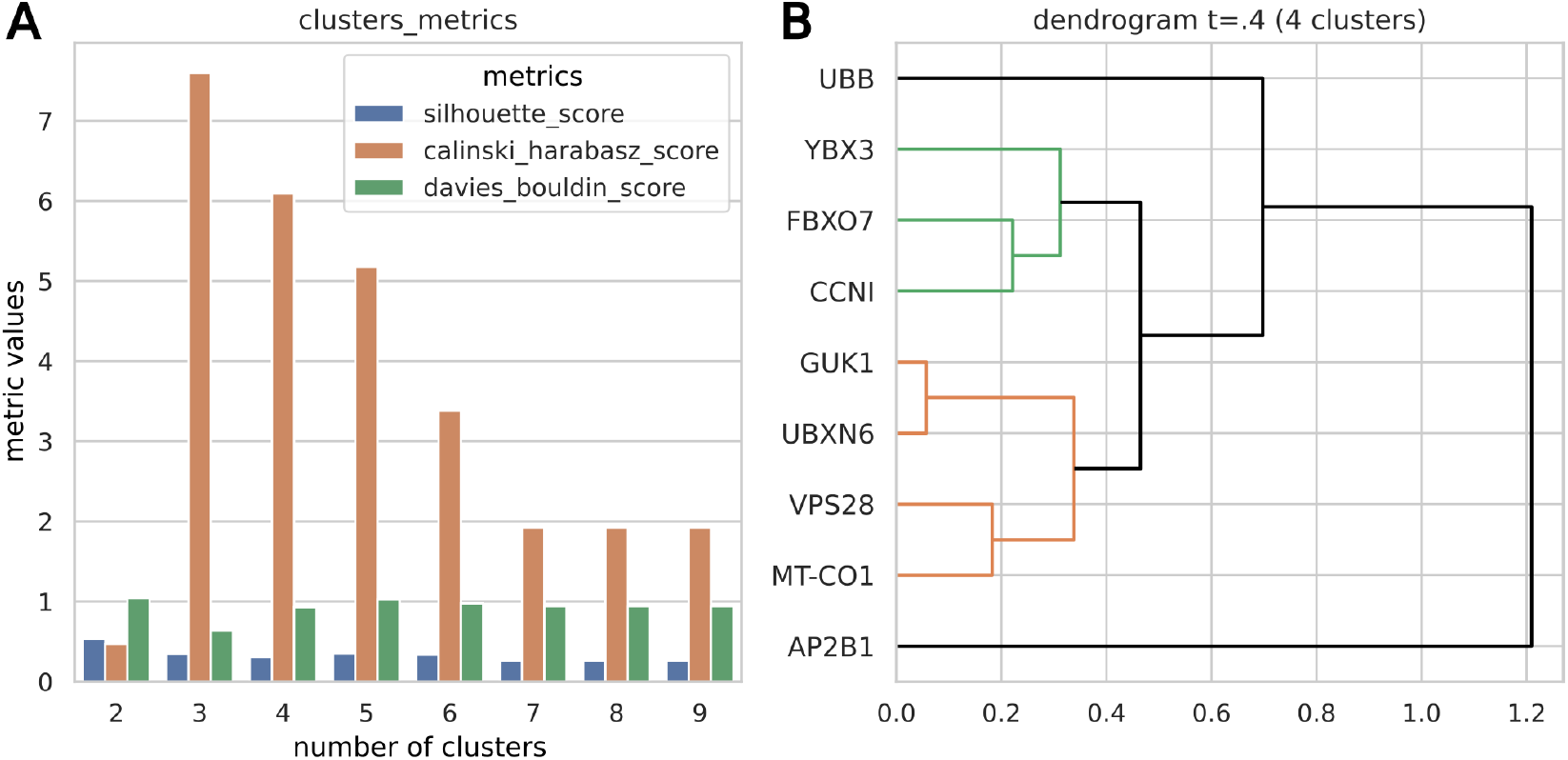
Hierarchical clustering to find groups that could be used as normalization factors. (A) The metrics results evaluate the optimal number of flat clusters to group HKGs based on correlation distance. The lower the silhouette score, the lower the Davies Bouldin score; the higher the Calinski Harabasz scores, the better the groups formed. We chose 4 clusters to define the groups since it is a reasonable number of groups and score metrics. (B) Average linkage hierarchical clustering based on correlation distance to visualize the groups formed.

**Fig. A4.**
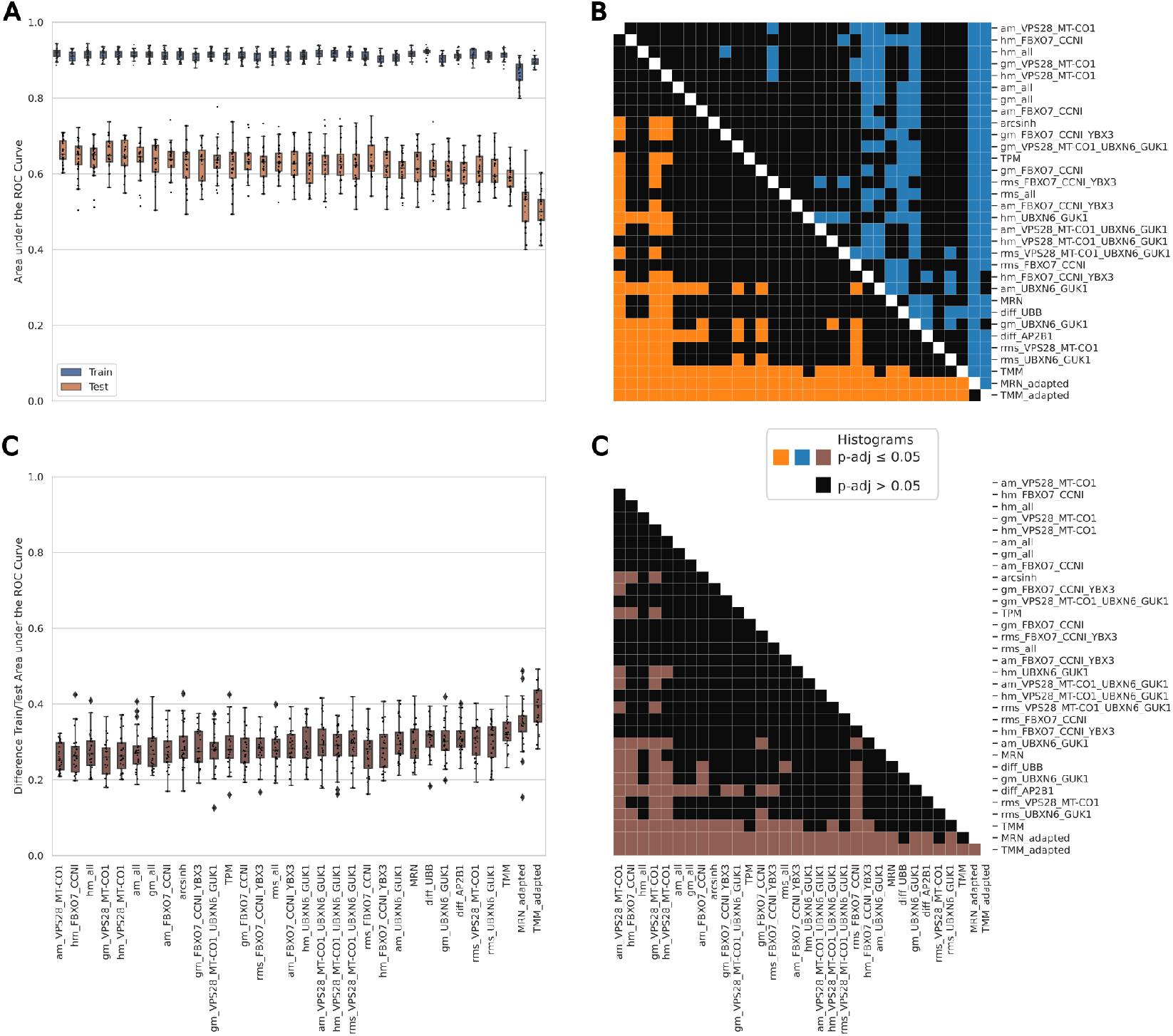
Analysis of data leakage in ALL normalization based on proposed housekeeping genes and benchmark methods. (A) Train and test values of AUC metric distributions for all different normalization methods for 25 cross-validations. To not inflate the adjusted p-values, we did this complete analysis after choosing the normalizations with higher AUC values (the arithmetic mean of genes VPS28 and MT-CO1, the harmonic mean of genes FBXO7 and CCNI, and the harmonic mean of all proposed HKGs. (C) The colored boxes represent statistically significant differences (adjusted p-value ≤ 0.05) in pairwise comparisons of AUC distributions between different normalization methods. The black box represents a non-significant difference. (D) Distributions of differences between train and test AUC values for each normalization method. (E) Adjusted p-values of pairwise comparisons between differences in AUC distributions for different normalization methods. The brown boxes represent statistically significant differences (adjusted p-value ≤ 0.05), while the black boxes represent a non-significant difference.

**Table A1.**
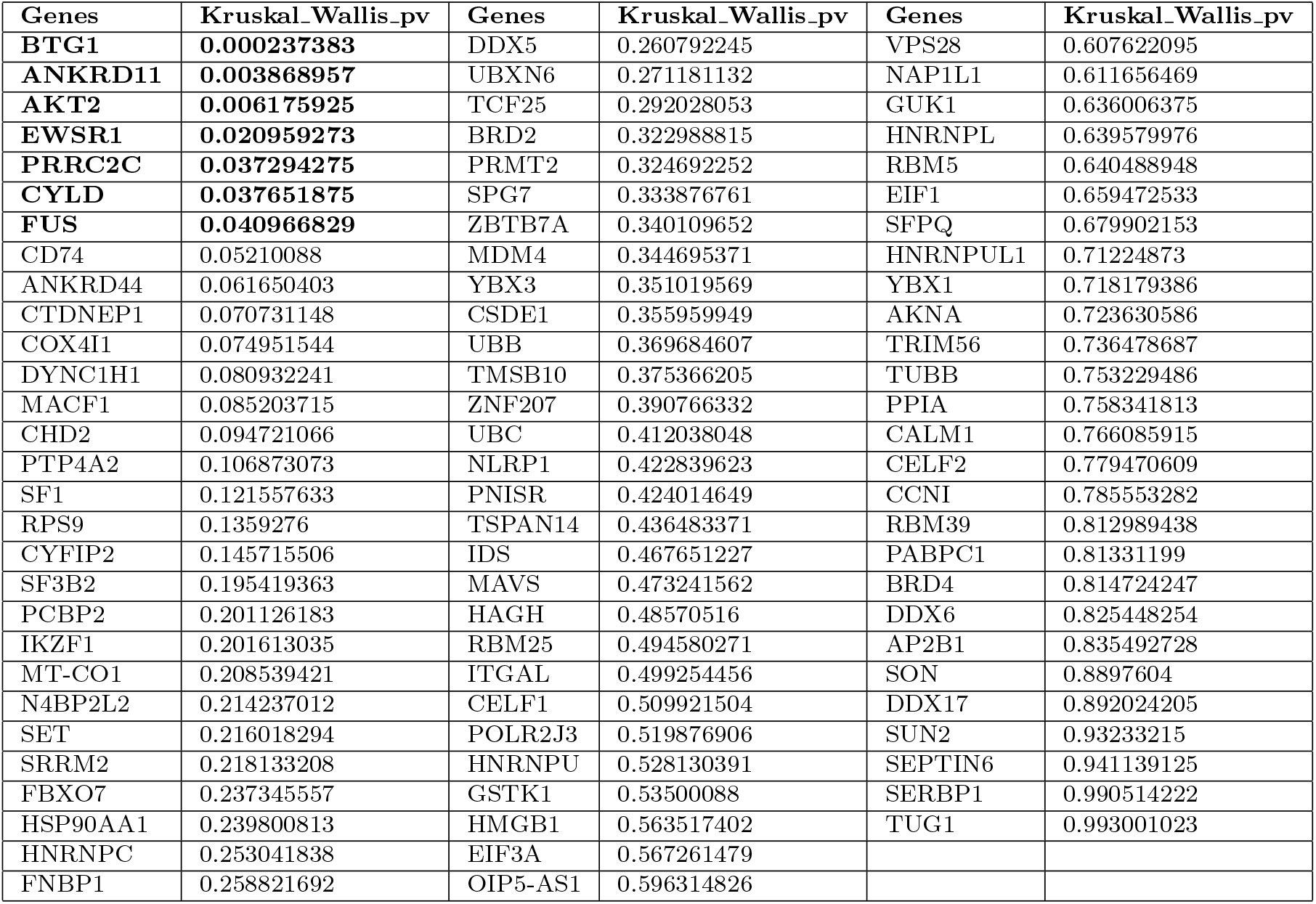
Kruskal-Wallis H test p-values. Test performed in dataset GSE175718 to verify if the expression distributions are different when comparing non-rejection, ABMR, and TCMR in transplanted patients. Statistically significant genes (p-value ≤ 0.05) are excluded as potential housekeeping genes; they are bolded in the table. We want to be conservative in exclusion, so we didn’t adjust the p-values. ABMR: antibody-mediated rejection; TCMR: T-cell-mediated rejection.

**Table A2.**
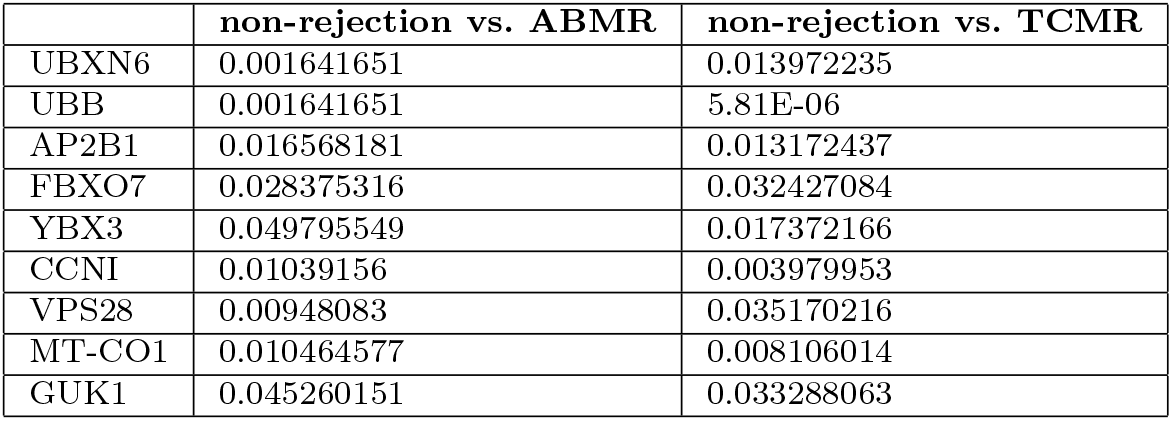
Two-One-Sided-Test with Brunner-Munzel test adjusted p-values. Performed to compare equivalence of expression distributions between non-rejection, ABMR, and TCMR with a Cohen’s d effect size at a maximum of 0.3. The significance level is adjusted p-value ≤ 0.05.

**Table A3.**
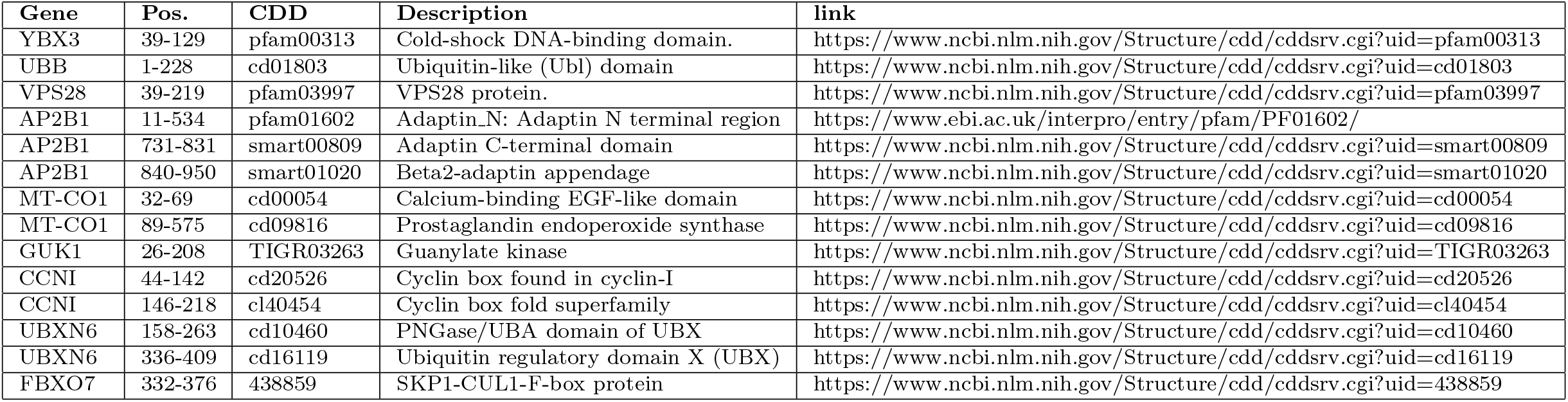
Important regions with low entropy in the HomoloGene and Conserved Domain Databases. Pos. is the position coordinate in protein. CDD is the code in the NCBI Conserved Domain Database. The links were accessed last time on 08/28/2024.

