## Supplementary material for "Housekeeping Gene Expression Normalization in Transcriptomics Mitigates Data Leakage in Machine Learning Models": Suplementary figures and tables

### Appendix A Supplementary Figures and Tabels

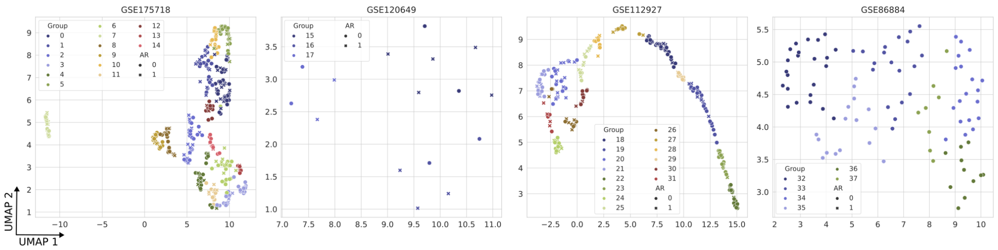


**Fig. A1 Unsupervised Louvain clustering to group samples due to lack of demographic information.** An unsupervised Louvain algorithm creates homogeneous groups for each RNA-seq dataset. The two dimensions of UMAP dimensionality reduction allow the visualizations of groups. The data from each cluster were used to calculate the coefficient of variation (CV), coefficient of variation of pairwise stability (cvSTB), and Gini coefficient.

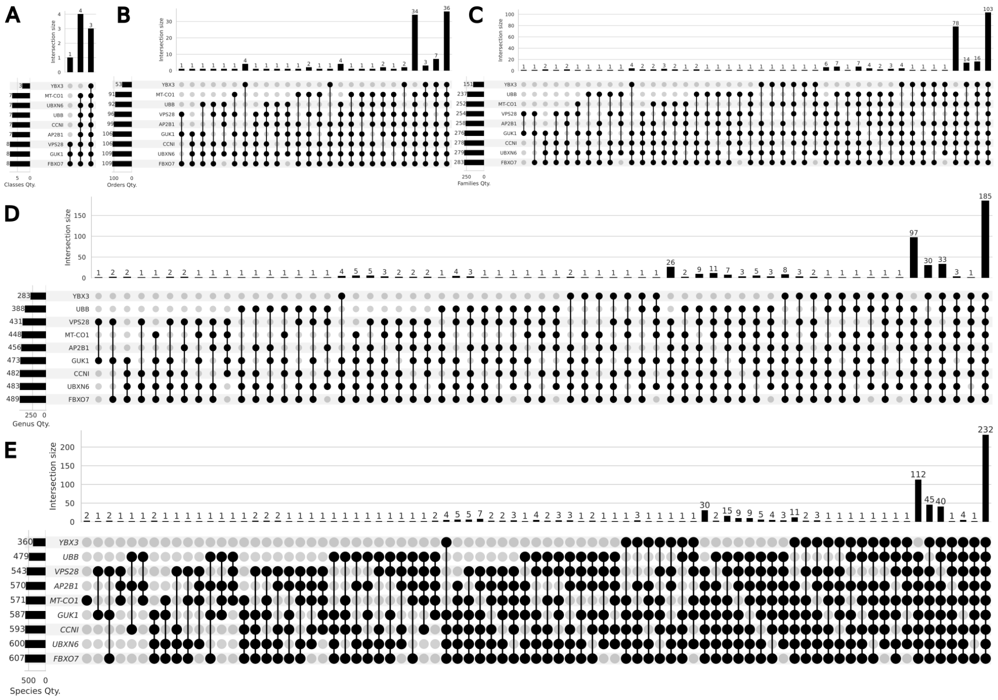


**Fig. A2 The upset plots with all combinations of homologs for Classes, Orders, Families, Genera, and Species.** (A) The upset plot of eight Classes that share homologs. (B) The upset plot of 111 Orders that share homologs. (C) The upset plot of 288 Families that share homologs. (D) The upset plot of 500 Genera that share homologs. (E) The upset plot of 630 Species that share homologs. The top barplot shows the number of species sharing the same homologous genes. The left barplot shows the total number of taxon with the homolog genes.

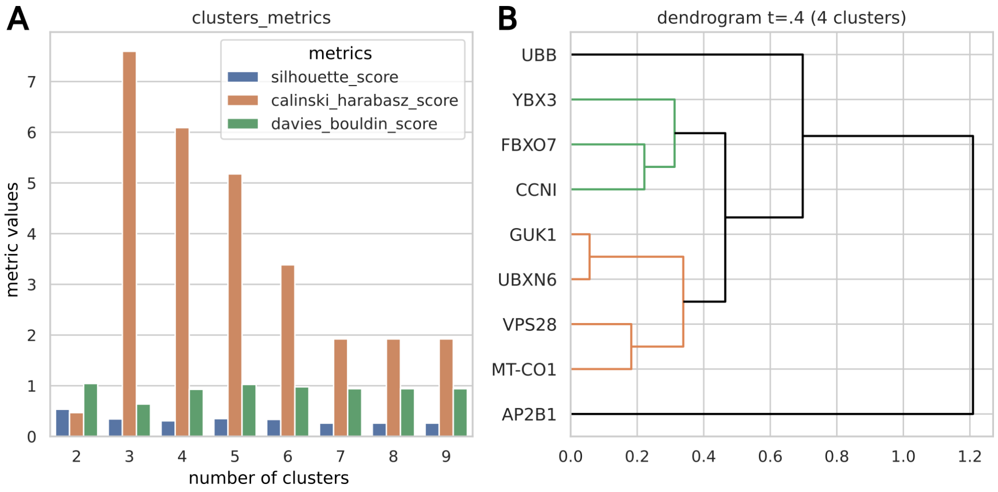


**Fig. A3 Hierarchical clustering to find groups that could be used as normalization factors.** (A) The metrics results evaluate the optimal number of flat clusters to group HKGs based on correlation distance. The lower the silhouette score, the lower the Davies Bouldin score; the higher the Calinski Harabasz scores, the better the groups formed. We chose 4 clusters to define the groups since it is a reasonable number of groups and score metrics. (B) Average linkage hierarchical clustering based on correlation distance to visualize the groups formed.

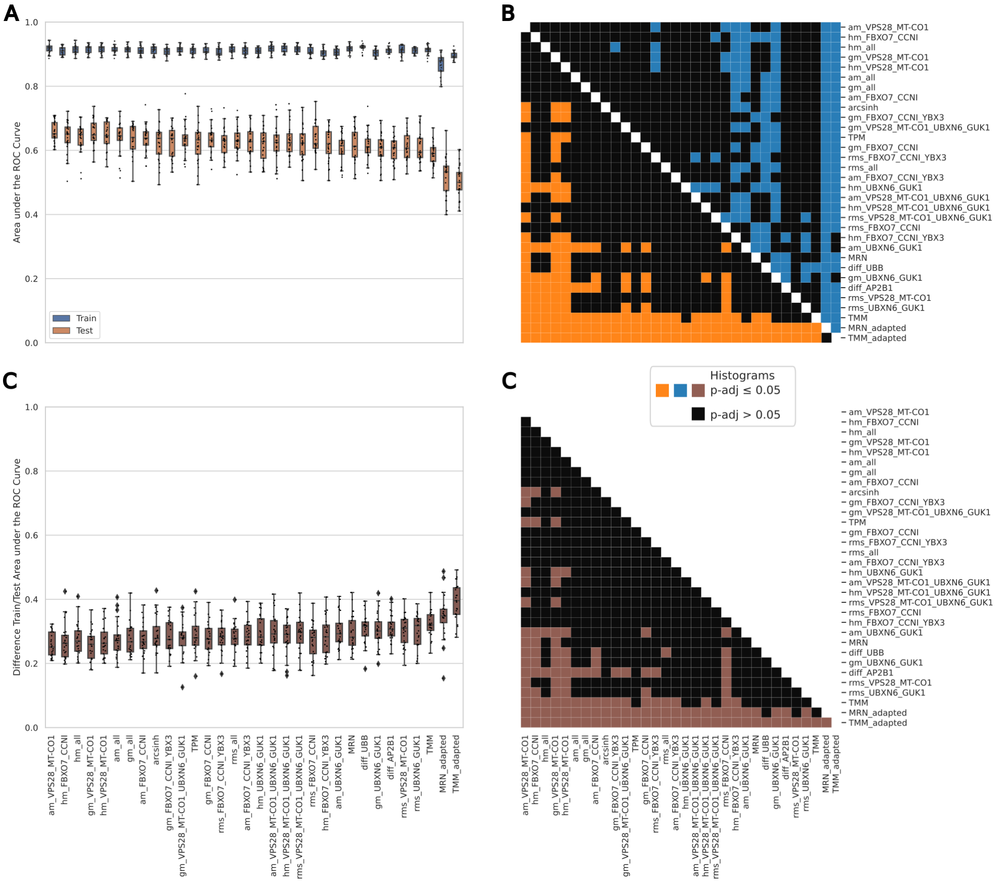


**Fig. A4 Analysis of data leakage in ALL normalization based on proposed housekeeping genes and benchmark methods.** (A) Train and test values of AUC metric distributions for all different normalization methods for 25 cross-validations. To not inflate the adjusted p-values, we did this complete analysis after choosing the normalizations with higher AUC values (the arithmetic mean of genes VPS28 and MT-CO1, the harmonic mean of genes FBXO7 and CCNI, and the harmonic mean of all proposed HKGs. (C) The colored boxes represent statistically significant differences (adjusted p-value $\leq 0.05$) in pairwise comparisons of AUC distributions between different normalization methods. The black box represents a non-significant difference. (D) Distributions of differences between train and test AUC values for each normalization method. (E) Adjusted p-values of pairwise comparisons between differences in AUC distributions for different normalization methods. The brown boxes represent statistically significant differences (adjusted p-value $\leq 0.05$), while the black boxes represent a non-significant difference.

**Table A1 Kruskal-Wallis H test p-values.** Test performed in dataset GSE175718 to verify if the expression distributions are different when comparing non-rejection, ABMR, and TCMR in transplanted patients. Statistically significant genes (p-value $\leq 0.05$) are excluded as potential housekeeping genes; they are bolded in the table. We want to be conservative in exclusion, so we didn't adjust the p-values. ABMR: antibody-mediated rejection; TCMR: T-cell-mediated rejection.

| Genes | Kruskal_Wallis_pv | Genes | Kruskal_Wallis_pv | Genes | Kruskal_Wallis_pv |
|---|---|---|---|---|---|
| **BTG1** | **0.000237383** | DDX5 | 0.260792245 | VPS28 | 0.607622095 |
| **ANKRD11** | **0.003868957** | UBXN6 | 0.271181132 | NAP1L1 | 0.611656469 |
| **AKT2** | **0.006175925** | TCF25 | 0.292028053 | GUK1 | 0.636006375 |
| **EWSR1** | **0.020959273** | BRD2 | 0.322988815 | HNRNPL | 0.639579976 |
| **PRRC2C** | **0.037294275** | PRMT2 | 0.324692252 | RBM5 | 0.640488948 |
| **CYLD** | **0.037651875** | SPG7 | 0.333876761 | EIF1 | 0.659472533 |
| **FUS** | **0.040966829** | ZBTB7A | 0.340109652 | SFPQ | 0.679902153 |
| CD74 | 0.05210088 | MDM4 | 0.344695371 | HNRNPUL1 | 0.71224873 |
| ANKRD44 | 0.061650403 | YBX3 | 0.351019569 | YBX1 | 0.718179386 |
| CTDNEP1 | 0.070731148 | CSDE1 | 0.355959949 | AKNA | 0.723630586 |
| COX4I1 | 0.074951544 | UBB | 0.369684607 | TRIM56 | 0.736478687 |
| DYNC1H1 | 0.080932241 | TMSB10 | 0.375366205 | TUBB | 0.753229486 |
| MACF1 | 0.085203715 | ZNF207 | 0.390766332 | PPIA | 0.758341813 |
| CHD2 | 0.094721066 | UBC | 0.412038048 | CALM1 | 0.766085915 |
| PTP4A2 | 0.106873073 | NLRP1 | 0.422839623 | CELF2 | 0.779470609 |
| SF1 | 0.121557633 | PNISR | 0.424014649 | CCNI | 0.785553282 |
| RPS9 | 0.1359276 | TSPAN14 | 0.436483371 | RBM39 | 0.812989438 |
| CYFIP2 | 0.145715506 | IDS | 0.467651227 | PABPC1 | 0.81331199 |
| SF3B2 | 0.195419363 | MAVS | 0.473241562 | BRD4 | 0.814724247 |
| PCBP2 | 0.201126183 | HAGH | 0.48570516 | DDX6 | 0.825448254 |
| IKZF1 | 0.201613035 | RBM25 | 0.494580271 | AP2B1 | 0.835492728 |
| MT-CO1 | 0.208539421 | ITGAL | 0.499254456 | SON | 0.8897604 |
| N4BP2L2 | 0.214237012 | CELF1 | 0.509921504 | DDX17 | 0.892024205 |
| SET | 0.216018294 | POLR2J3 | 0.519876906 | SUN2 | 0.93233215 |
| SRRM2 | 0.218133208 | HNRNPU | 0.528130391 | SEPTIN6 | 0.941139125 |
| FBXO7 | 0.237345557 | GSTK1 | 0.53500088 | SERBP1 | 0.990514222 |
| HSP90AA1 | 0.239800813 | HMGB1 | 0.563517402 | TUG1 | 0.993001023 |
| HNRNPC | 0.253041838 | EIF3A | 0.567261479 | | |
| FNBP1 | 0.258821692 | OIP5-AS1 | 0.596314826 | | |

**Table A2 Two-One-Sided-Test with Brunner-Munzel test adjusted p-values.** Performed to compare equivalence of expression distributions between non-rejection, ABMR, and TCMR with a Cohen's d effect size at a maximum of 0.3. The significance level is adjusted p-value $\leq$ 0.05.

| | non-rejection vs. ABMR | non-rejection vs. TCMR |
|---|---|---|
| UBXN6 | 0.001641651 | 0.013972235 |
| UBB | 0.001641651 | 5.81E-06 |
| AP2B1 | 0.016568181 | 0.013172437 |
| FBXO7 | 0.028375316 | 0.032427084 |
| YBX3 | 0.049795549 | 0.017372166 |
| CCNI | 0.01039156 | 0.003979953 |
| VPS28 | 0.00948083 | 0.035170216 |
| MT-CO1 | 0.010464577 | 0.008106014 |
| GUK1 | 0.045260151 | 0.033288063 |

**Table A3 Important regions with low entropy in the HomoloGene and Conserved Domain Databases.** Pos. is the position coordinate in protein. CDD is the code in the NCBI Conserved Domain Database. The links were accessed last time on 08/28/2024.

| **Gene** | **Pos.** | **CDD** | **Description** | **link** |
|---|---|---|---|---|
| YBX3 | 39-129 | pfam00313 | Cold-shock DNA-binding domain. | https://www.ncbi.nlm.nih.gov/Structure/cdd/cddsrv.cgi?uid=pfam00313 |
| UBB | 1-228 | cd01803 | Ubiquitin-like (Ubl) domain | https://www.ncbi.nlm.nih.gov/Structure/cdd/cddsrv.cgi?uid=cd01803 |
| VPS28 | 39-219 | pfam03997 | VPS28 protein. | https://www.ncbi.nlm.nih.gov/Structure/cdd/cddsrv.cgi?uid=pfam03997 |
| AP2B1 | 11-534 | pfam01602 | Adaptin_N: Adaptin N terminal region | https://www.ebi.ac.uk/interpro/entry/pfam/PF01602/ |
| AP2B1 | 731-831 | smart00809 | Adaptin C-terminal domain | https://www.ncbi.nlm.nih.gov/Structure/cdd/cddsrv.cgi?uid=smart00809 |
| AP2B1 | 840-950 | smart01020 | Beta2-adaptin appendage | https://www.ncbi.nlm.nih.gov/Structure/cdd/cddsrv.cgi?uid=smart01020 |
| MT-CO1 | 32-69 | cd00054 | Calcium-binding EGF-like domain | https://www.ncbi.nlm.nih.gov/Structure/cdd/cddsrv.cgi?uid=cd00054 |
| MT-CO1 | 89-575 | cd09816 | Prostaglandin endoperoxide synthase | https://www.ncbi.nlm.nih.gov/Structure/cdd/cddsrv.cgi?uid=cd09816 |
| GUK1 | 26-208 | TIGR03263 | Guanylate kinase | https://www.ncbi.nlm.nih.gov/Structure/cdd/cddsrv.cgi?uid=TIGR03263 |
| CCNI | 44-142 | cd20526 | Cyclin box found in cyclin-I | https://www.ncbi.nlm.nih.gov/Structure/cdd/cddsrv.cgi?uid=cd20526 |
| CCNI | 146-218 | cl40454 | Cyclin box fold superfamily | https://www.ncbi.nlm.nih.gov/Structure/cdd/cddsrv.cgi?uid=cl40454 |
| UBXN6 | 158-263 | cd10460 | PNGase/UBA domain of UBX | https://www.ncbi.nlm.nih.gov/Structure/cdd/cddsrv.cgi?uid=cd10460 |
| UBXN6 | 336-409 | cd16119 | Ubiquitin regulatory domain X (UBX) | https://www.ncbi.nlm.nih.gov/Structure/cdd/cddsrv.cgi?uid=cd16119 |
| FBXO7 | 332-376 | 438859 | SKP1-CUL1-F-box protein | https://www.ncbi.nlm.nih.gov/Structure/cdd/cddsrv.cgi?uid=438859 |
